# Unexpected *XIST* Expression in Male Hearts Associates with Disease

**DOI:** 10.1101/2025.04.09.648005

**Authors:** Gennady Gorin, Natalie DeForest, Linda Goodman

## Abstract

*XIST* is a long non-coding RNA that mediates the process of X chromosome inactivation in females, and has not been previously observed in males outside of developmental, pathological, and ectopic contexts. We report the unprecedented endogenous expression of *XIST* in 226 male heart samples across 15 single-cell RNA sequencing studies. The male expression of *XIST* is specific to a rare cell type typically annotated as neurons or glia. We investigate a variety of explanations and cross-reference the expression of *XIST* against age, sex, and pathology, finding signatures of a non-canonical inhibitory program, potentially mediated by a truncated transcript. Interestingly, the highest-powered datasets exhibit *XIST* upregulation in heart disease and comorbidity donors relative to healthier controls. Furthermore, in humans we identify a male-specific association of the *XIST* genomic locus with myocardial infarction (*p* = 1.3 *×* 10*^−^*^4^). Taken together, our findings suggest that the pathology of cardiomyopathy may involve uncharacterized, therapeutically actionable non-coding RNA pathways that operate through the cardiac nervous system, and underscore the importance of genomic investigations of the X chromosome.

## 1 Introduction

X-chromosome inactivation (XCI) establishes monoallelic expression of X-linked genes in female cells by condensing one of the X chromosomes. Although the details of the process are relatively complex and differ by species and tissue, several elementary points are generally considered uncontroversial:

- In humans, XCI is **ubiquitous** in and **specific** to females (XX) [1].
- XCI is mediated and continuously enforced by *XIST*, a long non-coding RNA (lncRNA) on the X chromosome [1].
- XCI leads to dramatic changes in the topology and accessibility of one of the copies of the X chromosome [2].
- Gene expression in the resulting Barr body (X_i_) ranges from near-total silencing to a slight decrease relative to the active X chromosome (X_a_) [3, 4], depending on the gene.
- There are rare exceptions to the ubiquity in females: in some restricted immune cell contexts, cells escape XCI [5, 6], leading to overexpression of X-linked genes.

Some rare exceptions to the specificity to females are more contentious. XCI and *XIST* expression transiently occur in the male (XY) germline [7–9]. However, *XIST* has also been reported in certain tumors, more tentatively in conditions with supernumerary sex chromosomes (e.g., with XXY or XXXY karyotypes) [7, 10], and recently claimed to be quantifiable by qPCR in male lymphocytes in a conference abstract, albeit without any follow-up reports [11]. *In vitro*, it has been observed in differentiating stem cells [12]. Ectopic induction in XY stem cells results in somewhat weaker XCI [13]. The expression of a truncated, non-silencing variant in male mice triggers autoimmunity to a *Xist* ribonucleoprotein complex [14]. In other words, there is some precedent for such exceptions, but they are well-understood to be atypical in somatic, differentiated tissues, and generally require ectopic expression.

The genomics and health implications of *XIST* have been almost exclusively studied in the context of XCI. Outside the recently reported non-canonical connections to autoimmunity [14], *XIST* and XCI escapees have been tentatively implicated in affective disorders [15] and aging [16]. Conversely, the induction of XCI has been proposed as a promising therapeutic approach to silence lethal trisomies [17]. However, the X chromosome and non-coding RNA are rarely included in genome-wide association studies [18–21], and remain an untapped reservoir of variation related to disease.

Separately, a great deal of effort has been expended toward characterizing the single-cell transcriptomes of human cardiac cells in healthy and pathological tissues [22]. At this time, there are several dozen such published datasets. Despite the variety of study designs and goals, the cell type compositions tend to be consistent with the “canonical” 2020 Human Cell Atlas dataset by Litviňuková et al. [23], used in lieu of a control in a number of studies [24–26]: cardiac tissue contains cardiomyocytes, vascular cells, fibroblasts, immune cells, adipocytes, and neurons. As in other tissues, all of these cell types can be sampled by single-nucleus sequencing, although adipocytes and myocytes are generally (but not universally [27]) considered morphologically incompatible with droplet-based single-cell protocols [23].

As the cardiac nervous system controls signaling and conduction, cardiac neurons have been investigated as potentially impactful and accessible targets for disease intervention long before single-cell sequencing. However, the architecture, role, and pathology of the heart’s neural population remains poorly characterized [28–30]. Analogously, in the single-cell atlases, the cardiac neurons are slightly more contentious than other cell types. The relevant cell population strongly expresses neurexins (*NRXN1* /*NRXN3*), proteolipid protein 1 (*PLP1*), myelin protein zero (*MPZ*), and a number of other nervous system markers [22]. The original heart cell atlas reported them as neurons, but noted that a subpopulation more closely resembles (glial) Schwann cells [23]. A follow-up study by the same authors remarked that the cells fail to express canonical neuronal markers and redefined them as glia [31]. This study additionally used staining to validate the presence of this distinct PLP1-expressing cell population and characterize its association with P cells. Similarly, spatial transcriptomics was performed to characterize the expression of the Schwann cell marker *MPZ* in a more recent study [32]. Other articles have reported these cells as neurons [25, 33, 34] or glia/Schwann cells [35, 36] with little further comment. Nominally, these are considered “prototypic” peripheral nervous system cells [22]. However, to acknowledge the remaining uncertainty in the field, we term this cell population “pseudo-glial.”

## 2 Results

### 2.1 Male pseudo-glia express *XIST*

Somewhat surprisingly, these pseudo-glia demonstrate high and specific expression of the *XIST* transcript in males (Fig 1). Although some variability is evident, the usual difference between male and female expression of *XIST* is 2–3 orders of magnitude: female samples show strong *XIST* expression in over half of all cells, almost certainly an underestimate due to experimental limitations; on the other hand, the expression in male samples is, although nonzero, vanishingly sparse and well within error bounds generally explainable by contamination, doublets, or barcoding errors. For instance, *XIST* expressors make up 63% of cells in female and 0.3% in male samples from the Litviňuková atlas [23]. Yet in the pseudo-glia, male and female cells have comparable amounts of *XIST*, with the latter showing slightly but not drastically higher expression. In fact, 67% of the rare *XIST* -positive male cells in the atlas are annotated as neural, and another 16% is assigned to an “unknown” population that strongly expresses the Schwann cell markers *PLP1* and *MPZ*. Overall, we observe average expression values that exceed 10*^−^*^2^ in 226 of the 248 male samples across fifteen studies of the human heart.

**Fig 1.**
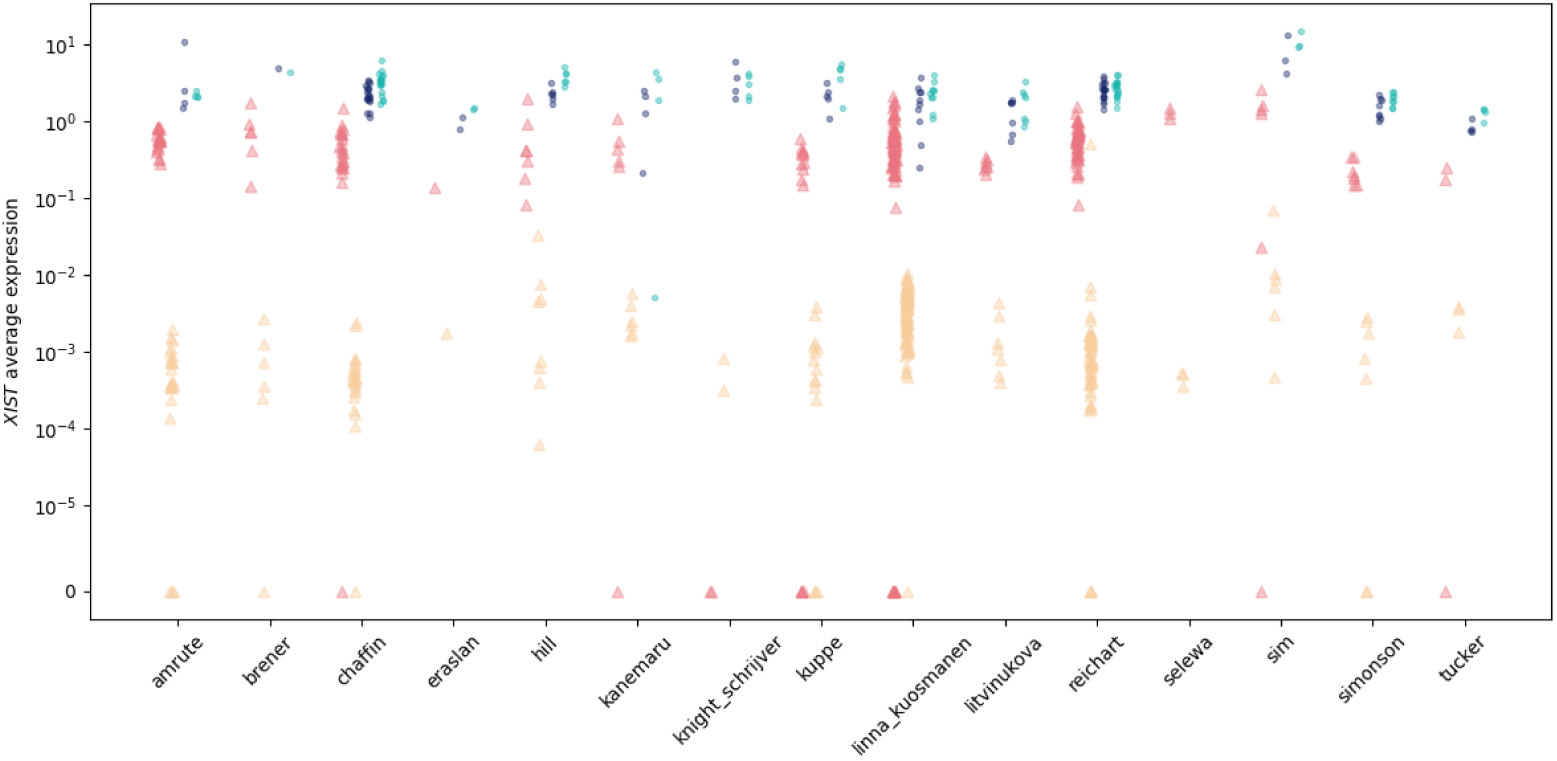
Male cardiac pseudo-glia show striking expression of the female-specific transcript *XIST*. Each point denotes average expression of the gene in one sample (typically one unique donor). Triangles denote samples reported as male. Circles denote samples reported as female. Red: male, pseudo-glial (cells annotated as Schwann cells, neurons, or glia). Yellow: male, all others. Dark blue: female, pseudo-glial. Teal: female, all others. To accommodate zero values, the region [0, 10*^−^*^5^] is linear.

### 2.2 Exogenous or artifactual explanations are inadequate

As expression of *XIST* in male somatic tissues is unusual, it is necessary to rule out potential exogenous sources. These include donor metadata errors, sample metadata errors, background contamination, doublets, and alignment errors.

Donor metadata errors refer to a tissue donor’s sex being reported incorrectly. This explanation cannot account for cell type-specific enrichment of *XIST*, as the transcript would be expressed at high levels elsewhere, as in other female samples. In addition, this type of error is fairly easy to spot: for instance, the donor AV3 in Kanemaru et al. [31], reported as a Caucasian female in her early 60s, expresses negligible *XIST* outside the pseudo-glia, but shows substantial expression of Y chromosome transcripts *UTY* (65% of cells), *DDX3Y* (43%), and *ZFY* (32%). This sample is readily evident in Fig 1, and displays the atypical high expression of *UTY* in Fig 2a. Unrecorded sex chromosome abnormalities, if they occurred, would be readily detectable by tissue-wide expression of *XIST* and Y-linked genes [7, 37, 38]. It appears most reasonable to conclude that the hundreds of donors under consideration, all of whom are annotated as male, express Y-linked genes, and do not express *XIST* outside the pseudo-glia, are indeed male (XY).

**Fig 2.**
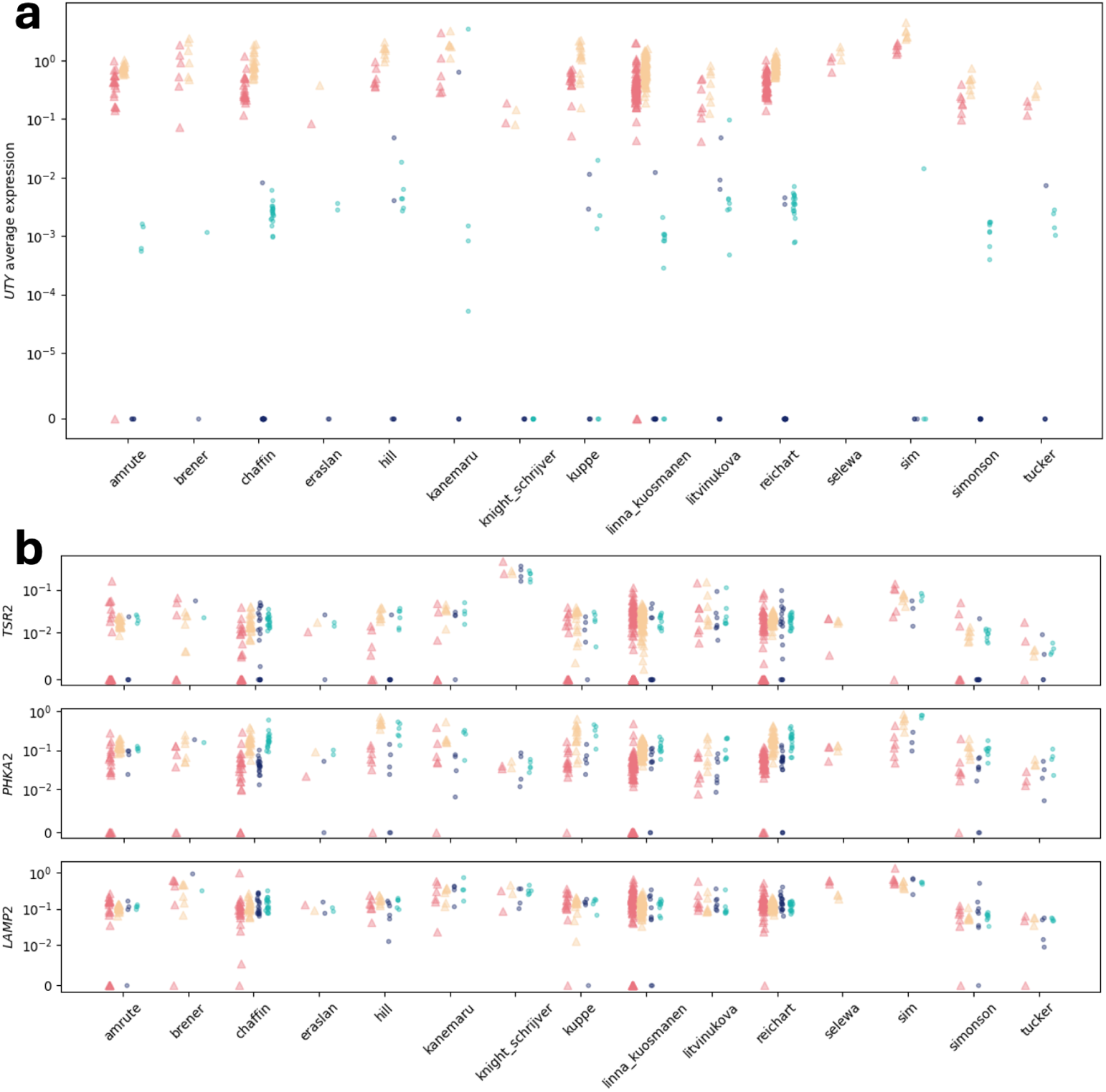
Expression of sex-associated genes in the heart. Conventions as in Fig 1. **a.** Average expression of *UTY* in cardiac datasets (linear region [0, 10*^−^*^5^]). **b.** Average expression of example genes reported [39] as consistently silenced in XCI by Tukiainen et al. (linear region [0, 10*^−^*^2^]).

Sample metadata errors refer to the relationship between a sequencing sample and a donor being reported incorrectly. A single sequencing run may contain several multiplexed donor samples, or tissues from single donor may have been sequenced separately, e.g. split by cardiac chamber, and the appropriate mapping may have been lost. Again, this explanation cannot account for the enrichment of *XIST* in a rare cell type and nowhere else. In addition, this type of error is also fairly easy to spot. For instance, in the Litviňuková atlas, donor H5, reported as a Caucasian female in her early 50s [23], has a relatively high expression of *UTY* (Fig 2a) exclusive to the right ventricular sample H0015_RV, which also has negligible *XIST* ; the other five samples assigned to this donor (LV, RA, LA, apex, and septum) have more typically female transcriptomes (high *XIST* and negligible Y-linked transcripts). In another example, the Reichart study reports that donor H40 is a male in his 60s [40]; the left ventricular sample BO_H40_LVW_premrna, one of four for this donor, shows a female-like transcriptome, where the others – two LV and one septal – show entirely typical male expression of *XIST* and Y-linked genes. Thus, metadata errors do not appear to be responsible for this effect.

Background contamination and doublets (e.g., due to incorporation of exogenous *XIST* from female samples processed in parallel) are implausible explanations, for a variety of reasons: the robustness of the expression across many studies, the presence of the same expression pattern in the male-only data by Selewa et al. [41], and the lack of a plausible mechanism for background transcript enrichment in a specific cell type. In other words, there is no clear source for exogenous *XIST*, nor a clear reason why it should specifically associate with pseudo-glia regardless of protocol.

Alignment errors are somewhat challenging to rule out. It is conceivable that a rare, cell type-specific transcript, generated from a different gene and unattested until now, shares enough structure to be mistakenly quantified as *XIST*. Such an explanation, appealing to genomic “dark matter,” is somewhat implausible if only because it would predict the total amount of “*XIST*” should be *higher* in female glia-like samples, rather than equal to or slightly lower than in other female cells, as actually observed in sequencing data. We do, however, note that observed data may be consistent with an uncharacterized *XIST* -like gene whose expression is cell type- and male-specific. Incidentally, overlap with *known* transcript sequences would artificially deflate “*XIST*” quantification. This is observed, e.g., in the Commons Cell Atlas reanalysis of the Litviňuková data (accession ERX4319109, sample H0026_RV, lane 3), which reports zero *XIST* expression [42, 43]. This loss of signal appears to be a consequence of an unstranded pseudoalignment strategy, which typically maps more reads, but cannot uniquely assign, and must discard, reads that can originate from either gene in a sense–antisense pair (i.e., exon 6 of the canonical transcript *XIST* -204, which overlaps *TSIX*).

The identification of the precise transcript identity requires investigation of the raw read alignments. None of the datasets reanalyzed in Fig 1 are accompanied by such alignments. However, a small unrelated study by Larson and Chin [44] released single-nucleus alignment data generated from the interventricular septa (IVS) of two males and two females. The reads aligned to genes in the *XIST* /*TSIX* region are displayed in Fig 7a (Section 4.3). The coverage in the male sample is considerably lower, but the localization appears to be consistent: reads are predominantly found in exon 6 of the canonical transcript, with far sparser expression in exon 1. Although these short-read data do not provide unequivocal evidence that the *XIST* transcript observed in human heart pseudo-glia is canonical, they do suggest that the count data are not likely to be attributable to, e.g., a poorly characterized unrelated transcript that happens to be localized to the same region, such as the pseudogene *LNX2BP*. These results agree with our reanalysis of a recent long-read single-cell heart dataset [45], which shows that pseudo-glial *XIST* read coverage is exclusive to exon 6 of the canonical transcript (Fig 7b, Section 4.4), biased toward the 3’ end and internal poly(A)-rich regions. The read coverage distribution is consistent with the relevant species being a truncated transcript that covers exon 6 of *XIST* -204, but we cannot rule out a more complex structure without full-length sequencing. In sum, there does not appear to be a plausible artifactual explanation for the expression of *XIST* in male hearts.

### 2.3 Cross-species context and biological implications

X chromosome inactivation is ubiquitous in mammals; in placental mammals, XCI is achieved through the expression of *XIST* [46, 47], although the transcript structure shows nontrivial variability [48]. Therefore, non-human mammals provide a natural point of comparison, as well as a potential choice of model organisms for follow-up *in vivo* investigations. However, the presence of *XIST* in non-human male pseudo-glia is ambiguous. As we discuss in Section 4.1.2, the absence of the lncRNA in genome references poses a barrier. Our preliminary investigation suggests that, in males, *Xist* is not expressed in the adult rat, mouse, pig, or cynomolgus monkey hearts, but shows some non-specific expression in the neonatal mouse heart. It is likely that other public datasets could shed light on the issue upon exhaustive reanalysis, but not so readily as the immediately evident signal in human tissues.

Despite the usual identification of *XIST* expression with XCI, there does not appear to be any evidence for canonical XCI in this cell population. Genes such as *TSR2*, *PHKA2*, and *LAMP2*, reported to be robustly inactivated in female tissues, appear to be no less expressed in male pseudo-glial cells than in others [39] (Fig 2b). Were the XCI process effective, one would expect their expression in XY cells to be far lower than in XX cells. However, this does not appear to occur; in fact, the non-pseudoautosomal (i.e., susceptible to XCI) X-linked gene *PLP1* is one of the strongest markers for this cell type. The inability to induce XCI is, to some extent, consistent with the results reported by Gayen et al. in an ectopic expression scenario [13]. This study’s results suggest that the inactivation mechanism relies on a ploidy counting mechanism mediated by escape products, and inactivation only occurs when the cell has two X chromosomes. However, despite a weaker magnitude, qualitative signatures of “canonical” silencing, such as *XIST* coating and silencing of X-linked genes, were observed in XY cells in that study. As the biological context is drastically different and the characteristics of *XIST* coating in the heart are unknown, we cannot establish whether the mechanisms proposed by Gayen et al. are relevant here.

The male expression of *XIST* does not appear to be restricted to these datasets, nor the heart. Similar male neurexin- and *XIST* -expressing populations are present in arterial atlases [49–51], albeit reported as smooth muscle cells or lymphocytes [49, 51]. Single-cell surveys of the human eye [52], musculature [53], prostate, and esophagus [36] report male donor *XIST* expression in rare Schwann cell populations, with some caveats (Section 4.1.1). In sum, the expression of *XIST* in male glia-like cells may not be tissue-specific; however, further investigation is hampered by the rarity of the cell type and the availability of data.

It is interesting to note that the fetal “neuronal” cell type reported by Knight-Schrijver [25] does not exhibit *XIST* expression in the two fetal male samples. This is consistent with the unusually low expression in the two fetal male samples reported by Sim et al. [54] (SAMN15889155 and SAMN15889156, visible in Fig 1). Aside from this dearth of expression in developing heart tissues, the expression of *XIST* is remarkably stable, varying across approximately an order of magnitude across the entire human lifespan. This consistency is more striking yet due to the small sample sizes and the lack of adjustments for technology, batch effects, pathology, or discrepancies in reference annotations (Fig 6a), all of which should be expected to introduce variability. After accounting for age and source study as covariates, the age effects on expression, if any, appear to be minimal (Section 4.2.4).

To investigate the transcriptome-wide associations of *XIST* expression while accounting for the many technical and biological covariates, we performed differential expression (DE) testing between *XIST* -positive and *XIST* -negative cells annotated as neurons or glia, encoding the donor ID as a model covariate (Section 4.2.3). As shown in Fig 3, the *XIST* -positive cells exhibited a striking decrease in a diverse variety of genes in male samples, and little to no difference from the *XIST* -negative cells in female samples.

**Fig 3.**
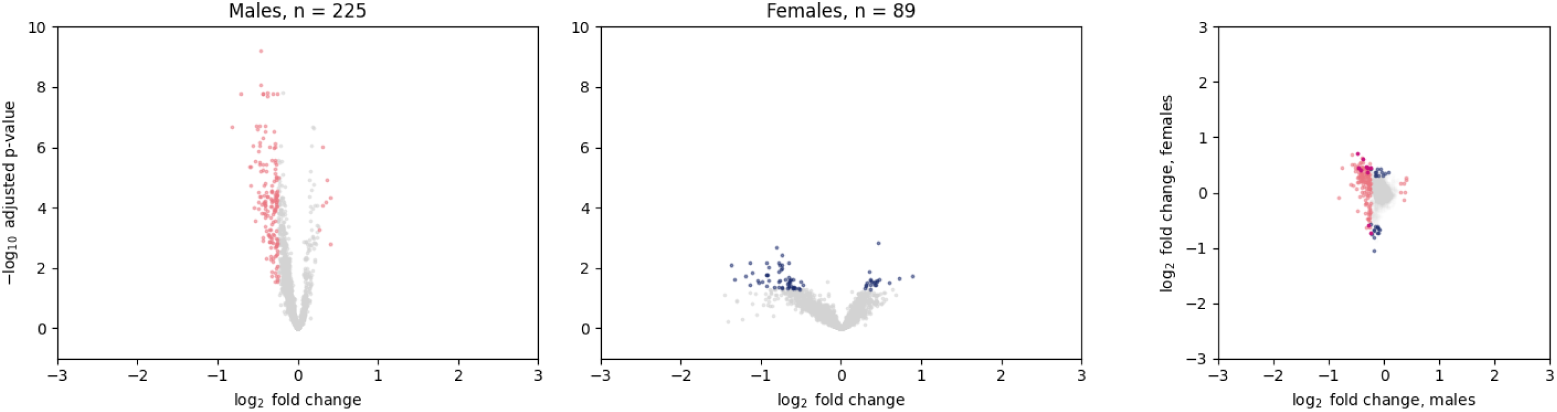
Results of differential expression testing between *XIST* -positive and *XIST* -negative pseudo-glial cells. Left: male samples (red: DE genes). Middle: female samples (blue: DE genes). Right: relationship between log_2_ fold changes inferred for male and female samples (red: male DE genes; blue: female DE genes; pink: genes DE in both comparisons). Differential expression is defined as | log_2_ FC*| >* 0.25 and adjusted *p*-value *<* 0.05. *XIST* excluded from visualization. Visualization restricted to genes with baseMean *>* 5.

Gene Ontology (GO) analysis of the 225 genes differentially expressed in female samples reported few significant enrichments, primarily the cellular and mitochondrial ribosomal proteins. Restricting the analysis to the 75 genes with at least modest average expression (baseMean *>* 5) yielded analogous enrichment of cellular ribosomal proteins and electron transport chain components, with modest statistical significance. It is far from clear that even these minor differences are meaningful, as the distinction between *XIST* -positive and -negative cells in female samples is likely to be predominantly governed by incidental loss of molecules in library construction and sequencing, rather than true lack of *XIST*, which is understood to be ubiquitous in XX cells. This is supported by the lack of detectable signatures of, e.g., escape from XCI in supposedly *XIST* -negative cells. In contrast, XCI escape *is* readily detectable through analogous statistical testing between male and female donors (Section 4.2.2).

GO analysis of 167 DE genes in male samples (likewise restricted to baseMean *>* 5) reported a variety of enriched biological processes, including genes associated with myofiber assembly and myogenesis (various components and forms of titin, fibulin, myomesin, fibronectin, actinin, and actin), with action potential regulation (channels and receptors *SLC8A1*, *CACNB2*, *CACNA1C*, and *RYR2*, alongside *FGF12*), with neuron migration regulation (*PHACTR1*, *UNC5C*, *ERBB4*, and *NRG3*), among others.

These genes were predominantly downregulated; the few exceptions with higher expression in *XIST* -positive cells included several relatively sparsely observed genes, including the potassium and choline transporters *KCNMA1* and *SLC5A7*, the glutamate receptor *GRID2*, the immunoglobulin *OPCML*, the obscure sulfotransferase *CHST9* (tentatively associated with heart conditions in genomics studies [55–57]), the neuronal development mediator *ALK*, the receptor modulators *LYPD6* and *OPCML*, and the poorly characterized gene *FAM135B*. None of these genes are sex-linked. Three X-linked genes, all outside the pseudoautosomal regions, were downregulated in *XIST* -positive cells: *TMSB4X*, *ARHGAP6*, and *DIAPH2*. All are involved in actin binding.

The pervasive depletion of canonical muscle transcripts in *XIST* -positive cells is suggestive, although some caution is warranted: these gene products are ubiquitous in, e.g., the cardiomyocytes, and a nontrivial fraction of the expression is likely exogenous due to contamination from the cell lysate. However, it is not clear how such a mechanism might specifically contaminate *XIST* -negative cells in male samples. We discuss these potential explanations in Section 4.2.3.

## 3 Conclusion

The potential biological impact of this finding is unclear. It is possible that the male *XIST* expression is incidental or explicable by artifacts not considered here, and the cells may be “glia as usual.” However, the fundamental divergence between these cells and supposedly universal patterns of gene expression regulation elsewhere in the human body leads us to urge caution in treating them as “prototypic” [22], as the functional differences may be far more profound. At the very least, the expression patterns reported here are striking counterexamples to the conventional wisdom that *XIST* is sufficient for XCI and that its expression is strictly pathological or developmental in males.

It is tempting to speculate that the suppression of the myogenesis program in the *XIST* -positive population is effected by a direct mechanism analogous to XCI. It is also likely mistaken: the suppression is genome-wide and seemingly specific, rather than localized to a single chromosome. It appears more plausible that *XIST* is involved upstream to a set of regulators that induce the usual myogenesis program, which are suppressed as the result of its action on some narrow, and unknown, set of mediators. The ultimate physiologic purpose is not immediately clear, although the context of the niche may provide a clue: the extracellular environment of the heart is likely to be awash in endocrine and paracrine signals that induce myogenesis; these signals may well infiltrate non-muscular cells and induce this transcriptional program; some baseline level of its expression may be generally tolerable, but interfere with the function of this particular cell type; *XIST* -mediated inhibition happens to be the mechanism of choice for suppressing it. However, the causal and molecular details are unknown: *XIST* expression may mediate the effect, or merely be an incidental consequence of establishing the pseudo-glial transcription program. Even if *XIST* plays an active role, the mechanism may involve the regulation of X-linked genes in *cis*, a non-canonical pathway in *trans* [58, 59], or an altogether new mechanism, possibly through an uncharacterized isoform. Overall, it is challenging to ascertain the role of the cell type or its transcriptional program based solely on bioinformatic data; experimental validation would be necessary, although the apparent lack of homology to model organisms presents a serious barrier. Nevertheless, a recent study [60] independently identified this expression trend, proposed that the relevant *XIST* species is a truncated transcript generated from an alternative promoter, and characterized the non-XCI effects of its overexpression in HEK-293T cells, suggesting the relevant mechanisms may be identifiable *in vitro*.

It is additionally tempting to connect this atypical cardiac expression of a sex-linked gene in males to the increased incidence of heart failure in males [61]. Based on the relatively scant evidence, the expression of *XIST* and the suppression of the reported program may plausibly be protective, deleterious, or both depending on the broader context of the organ and the organism. In Section 4.2.5, we examine this question, and find that the expression of *XIST* appears to be higher in samples with pathology or comorbidities in three relatively large datasets, but the smaller datasets are less illuminating.

In principle, this scenario should be an ideal application of genomics: the gene is specific to a single cell type, so any genome-wide association study (GWAS) results connecting variability in male *XIST* to cardiac phenotypes should reflect the functional role of *XIST* in the pseudo-glia. Unfortunately, sex chromosomes are generally omitted from GWAS [18–20]. For instance, a recent GWAS of the determinants of dilated cardiomyopathy in tens of thousands of patients and over a million controls analyzes and reports some fifteen million variants, all on the autosomes [62]. The Cardiovascular Disease Knowledge Portal, which aggregates hundreds of genetic association datasets, reports one SNP (rs1474563) in the *XIST* region, associated with height [63]. The only pathogenic variant explicitly reported in dbSNP [64] is rs773396320, just upstream of the first exon of *XIST*, which leads to skewed XCI [65]. In addition, in spite of continuing efforts, variants in lncRNAs remain understudied [21]. Whether due to less study, greater exposure to natural selection [18], or selection biases (for instance, if *XIST* loss of function in males leads to embryonic mortality), the data are sparse and do not illuminate the cardiac role of *XIST*.

To enhance sensitivity to male-specific effects, we interrogated the *XIST* genomic locus in a sex-stratified association study for myocardial infarction [66]. In males, we identified a significant association of the *XIST* locus with myocardial infarction (*p* = 1.3 *×* 10*^−^*^4^, *n* =160,316 cases / 6,672 controls), defined by the variant rs187334110 for which *XIST* is the nearest gene. This association of *XIST* with myocardial infarction is not seen in females (*p* = 0.3, *n* =192,586 cases / 1,567 controls), suggesting that variation at this locus may contribute to heart disease in a sex-dependent manner. This connection is intriguing, and suggests that the dearth of large-scale X-chromosomal studies represents a lacuna that may reveal ties to sex-specific pathology under further investigation.

Alternatively, the role of *XIST* may be elucidated by cross-referencing its pseudo-glial expression against the male donors’ genotypes. For instance, a particular polymorphism may be associated with higher *XIST* expression in one of studies considered here, and with greater risk of cardiac conditions in the context of a larger GWAS. Several of these studies summarize the representation of known heart condition-associated variants in expression or chromatin accessibility data, albeit on the level of peaks or counts and without donor genotyping [31, 33, 41, 67]. To our knowledge, the only large study reporting single-cell data alongside patient genotypes is the investigation of pathogenic variants in dilated cardiomyopathy by Reichart et al. [40]. In this study, variant identification was predominantly performed through 46- or 174-gene Illumina TruSight sequencing panels. Nine male donors were genotyped through whole exome or whole genome sequencing; however, neither raw nor intermediate genotyping data were released alongside the controlled-access transcriptomic data (EGAS00001006374). Many studies associate bulk RNA expression with variants [68, 69], but drawing conclusions about cell type-specific expression from bulk data remains problematic [70]. Therefore, characterizing the genetic correlates of male *XIST* expression in an unbiased way does not seem feasible at this time.

This analysis is necessarily cursory. Ideally, such analysis should be performed on harmonized data, quantified using the same reference and conventions. For instance, each of the investigated datasets quantified between 24,250 and 39,464 features; 10,391 overlapped between all studies, and only those were used for DE gene identification. It is plausible that biologically interesting effects are masked by the incidental omission of one or another gene due to choice of reference. In addition, processing and cell type assignment are highly subjective and manual, and seemingly arbitrary choices of quality control thresholds or hyperparameters typically lead to drastic differences [71]. The pseudo-glial population investigated here is distinctive and the detection of *XIST* is robust, but takes existing annotations at face value, and may be more or less reproducible in a comprehensive, consistent reanalysis. Such reanalysis would make this study more systematic, but would necessarily be a manual task, as any tools automating the process would have been trained and tested on the very datasets under scrutiny. Naturally, the reanalysis of raw data is impractical — in most cases impossible without institutional approval due to patient privacy concerns — and would encounter the same issues related to ambiguous or contradictory metadata reported here. It is also immaterial to the key finding, which is clear even on a superficial and qualitative investigation of publicly available data.

Finally, this unusual finding in plain sight, revealed by lncRNA expression, echoes, and is directly inspired, by recent discovery of severe, previously uncharacterized artifacts in single-cell data, wherein nuclear transcripts are absent from subpopulations of certain cell types, perhaps suggesting quality control issues [72, 73]. We suggest that the biological result, as well as severe metadata errors in large studies, should urge stronger scrutiny of results, as well as a healthy amount of skepticism regarding approaches for automating analyses by summarizing the existing literature, or by tacitly accepting the results of potentially incomplete analyses as “ground truth.”

## 4 Methods

### 4.1 Dataset curation and analyses

#### 4.1.1 Human data acquisition

##### Amrute

The dataset released by Amrute et al. was acquired from GEO at GSE226314 [35]. The Seurat RDS file was converted to AnnData using the SeuratDisk R package. The orig.ident field was used as donor ID and cell.type field was used as cell type. All non-pathological donor samples were omitted, as they had been collected as part of an earlier study [74].

Sexes and ages were annotated manually based on metadata in Supplementary Table 1. Donors 388 and 378, listed as nonresponders in the dataset, were not listed in metadata, nor disclosed on the GEO entries. Sexes were inferred to be male based on expression of Y-chromosomal genes. Donors 370 and 371, listed as female and male nonresponders in the metadata, were not present in the dataset.

Pre- and post-LVAD sample IDs were retained in their original form, somewhat arbitrarily treating the time points as though they had been collected from distinct donors. This is a simplification, targeted at avoiding the elision of nontrivial remodeling effects reported by Amrute et al. It is possible that a more sophisticated statistical design would obviate the need for this simplification.

##### Brener

The count data for the dataset released by Brener et al. was acquired from GEO at GSE185457 [24]. These data do not have cell type annotations. The annotations were obtained from a version of the dataset previously available at the Broad Single Cell Portal (SCP1464), since removed from the database. Sexes and ages were annotated manually, based on metadata in Supplementary Table 5; donor 15_144548-RV was female and all others were male. Only COVID-19 samples, collected for this study, were retained. The batches field was used as donor ID and leiden field was used as cell type.

##### Chaffin

The AnnData object released by Chaffin et al. was acquired from the Broad Single Cell Portal (SCP1303) [75]. The donor_id field was used as donor ID and cell_type_leiden0.6 field was used as cell type. Ages were parsed using the age field.

##### Eraslan

Data were obtained from CZI CELL*×*GENE and filtered for the heart samples, which had the tissue annotation “anterior wall of left ventricle” [36]. Ages were annotated manually, based on metadata in Table S1; these data used different bins than the released AnnData file (suggesting, e.g., that the age of donor GTEX-1ICG6 was 70, as it is listed in the 70–79 bin in one source and the 61–70 bin in the other). The donor_id field was used as donor ID and cell_type field was used as cell type.

Somewhat surprisingly, male donor GTEX-1HSMQ exhibited high *XIST* expression outside the heart Schwann cells, albeit only in the lung tissue sample. This does not appear to be a sample metadata artifact, as *UTY* is also ubiquitous throughout. This expression pattern is somewhat challenging to account for.

##### Hill

Single-nucleus count data were obtained from the Broad Single Cell Portal (SCP1852) and merged into an AnnData structure [76]. The donor_id field was used as donor ID and MainCellType field was used as cell type. Ages were parsed using the age field, rounding down to the year. Donor 13_198, which had two distinctly named samples corresponding to left and right ventricular samples, was kept split. UK1 and UK2 samples were mapped to FC3CB and 3B62D based on manual cross-referencing of ages.

##### Kanemaru

Data were obtained from CZI CELL*×*GENE and subset to primary data [31]. One cell attributed to donor D11 was removed, yielding 211,060 cells, in agreement with Fig. 1 of the original publication. Ages were annotated manually, based on metadata in Supplementary Table 1; these values largely agreed with those reported in the released AnnData structure, with the exception of donor AH2, reported as 45–50 years old in the table but 40–45 in the data file. The donor_id field was used as donor ID and cell_type field was used as cell type.

##### Knight-Schrijver

Data were obtained from CZI CELL*×*GENE and subset to primary data [25]. As all primary data were fetal, ages were set to 0. The donor_id field was used as donor ID and cell_type field was used as cell type.

##### Kuppe

Data were obtained from CZI CELL*×*GENE [33]. Ages were parsed from the development_stage field. The donor_id field was used as donor ID and cell_type field was used as cell type.

##### Linna–Kuosmanen

Data were obtained from CZI CELL*×*GENE [32]. The two datasets (Periheart and Carebank) were concatenated. The donor_id field was used as donor ID and cell_type field was used as cell type. The number of donors was rather larger than that reported in the original article and its metadata. We did not filter the dataset further. Ages were parsed from the development_stage field.

##### Litviňuková

Data were obtained from CZI CELL*×*GENE [23]. The donor_id field was used as donor ID and cell_type field was used as cell type. Ages were annotated manually, based on metadata in Supplementary Table 1; the bins reported in the latter were finer than those disclosed in the dataset.

##### Reichart

Data were obtained from CZI CELL*×*GENE and subset to primary data [40]. The donor_id field was used as donor ID and cell_type field was used as cell type. Ages were parsed from the development_stage field using the metadata provided in Supplementary Table 1.

##### Selewa

Data were obtained from CZI CELL*×*GENE and subset to primary data [41]. The donor_id field was used as donor ID and cell_type field was used as cell type.

##### Sim

Data were obtained from CZI CELL*×*GENE and subset to primary data [54]. The donor_id field was used as donor ID and cell_type field was used as cell type. Ages were parsed from the development_stage field. All fetal samples’ ages were set to 0.

##### Simonson

Data were obtained from the Human Cell Atlas Data Explorer [77]. The donor_id field was used as donor ID and cell_type_leiden0.5 field was used as cell type.

##### Tucker

Data were obtained from the Broad Single Cell Portal (SCP498) [67]. The biological.individual field was used as donor ID and Cluster field was used as cell type. Ages and sexes were manually annotated based on Table 1.

#### 4.1.2 Non-human

There is no evidence for *Xist* expression in pseudo-glia outside the humans, and relatively little data to draw upon to confirm or disconfirm its presence, despite the publication of several nonhuman heart sequencing studies. In general, analogous cell populations are robustly identifiable, but *Xist* is either not observed or not included in the genome reference. In this section, we summarize potential comparable data sources.

A 2018 study by Skelly et al. reports Schwann cells in mouse heart and uses *Xist* and Y-linked gene expression to effectively demultiplex pooled male and female samples, reporting Schwann cell differences between sexes but not remarking on any detection of, e.g., cells simultaneously positive for *Xist* and *Uty*, making their presence dubious [78]. The 10X scRNA-seq dataset generated as part of the Tabula Muris Senis project reports 58 cardiac neurons from male subjects, with no *Xist* expression, but this sample size may be too small to be conclusive or representative [79]. Several other developmental datasets have been collected; however, they do not report glial or neuronal cell types [80–82].

A study of sex-specific pressure overload in cats [83] reported a “neuronal/conduction” cluster with prominent expression of *NRXN1* but not *PLP1* or *NRXN3*. However, this dataset did not contain *XIST* in the reference, although the presence of *XIST* in cats is well-known and often serves as the first pedagogical introduction to X inactivation, using the calico pattern as a case study [84]. Similarly, the PANDORA atlas of non-model species reports small *NRXN1* /*PLP1* -expressing clusters in the cat and tiger hearts, annotated as cardiomyocytes and fibroblasts respectively [85]. However, neither dataset contains *XIST* expression data, presumably due to reference limitations.

##### Mouse: Mesquita, Li, Patrick, and Ranjbarvaziri via scBaseCount

A recent initiative, scBaseCount, attempts to facilitate the consistent analysis of transcriptomic data by applying a common quantification procedure to a large cross-section of public 10X single-cell datasets and hosting the results [86]. Without aiming to be exhaustive, we acquired a diversity of adult mouse heart datasets: two samples from Mesquita et al. [87], five from Li et al. [88], six from Patrick et al. [89], and four from Ranjbarvaziri et al. [90]. All mice were male, as confirmed by the expression of Y-chromosomal markers. However, all had relatively low expression of *Xist* (6–33 UMIs per sample). Clustering the *scVI* embedding of the data yields some 2,000 cells substantially enriched in the pseudo-glial markers [91, 92], but 0–1 UMIs of *Xist* per sample, suggesting that the population is present, but does not express the gene. Therefore, there does not appear to be any convincing evidence suggesting that the expression of *Xist* occurs in adult mouse hearts.

##### Mouse: Shen

A small study characterizing the transcriptomic landscape of the neonatal mouse heart, using four donors from the first six days after birth, reported a very small *Plp1* -containing neuronal population [27, 93]. We acquired the count matrices, generated using Cell Ranger 5.0.0, from GSE232466. The study did not determine the pups’ sex and did not disclose the cell type assignments. However, inspection of the count matrices suggested that the pups collected on days 1, 2, and 4 were male, as evidenced by relatively high (*>* 200 UMIs) expression of the Y-chromosomal genes *Uty*, *Ddx3y*, and *Kdm5d* ; these genes showed negligible expression (*<* 5 UMIs) in the pup collected on day 6.

Interestingly, we observed remarkably high *Xist* expression (*>* 40, 000 UMIs) in all samples, with the highest expression (*≈* 150, 000 UMIs) in the putative female sample. These quantifications translate to average male expression of 3.5–6.5 UMIs per barcode. Clustering the *scVI* embedding of the data yields 127 cells substantially enriched in the pseudo-glial markers [91, 92]. The presence of *Xist* was ubiquitous, and not restricted to the neuronal subpopulation, which had a similar average *Xist* expression (0.6–6.5 UMIs per barcode). In sum, although the male neonatal mouse heart expresses *Xist* to a surprisingly large degree, it does not appear to do so in a cell type-specific fashion.

For the sake of completeness, we note that *Xist* was depleted in one cluster, which expressed canonical muscle marker genes (*Myl2*, *Myl3*, *Tnnt2*), but did not express *Malat1*, a ubiquitous nuclear transcript. The lack of nuclear content — noted by the original authors — strongly suggests that this cluster is artifactual, and represents the *technical* failure to lyse the cardiomyocyte nucleus rather than the detection of a novel physiological cell type, previously reported in other tissues [72, 73].

##### Rat: Arduini

A recent large study consisting entirely of male Wistar rats [94] reported analogous glia-like cell types, but was based on a bespoke version of the Ensembl *R. norvegicus* Rnor_6.0.96 genome without *Xist*. We realigned the raw data to quantify *Xist* expression.

We acquired raw reads for 19 left ventricular samples from GSE280111 [94]. We generated a pseudoalignment reference from the RefSeq GRCr8 genome (“genomic” GCF_036323735.1), using *kallisto | bustools* 0.28.2 [95]. We ran kb ref in the nac configuration, using the d-list option to add potential artifactual sequences — the 10X template-switch oligo, its reverse complement, and the (A)_30_ oligomer — to the D-list for masking. We pseudoaligned the left ventricular samples using kb count in the nac configuration and inspected the unfiltered total count matrices to characterize the *Xist* content of the sequencing libraries. The datasets had negligible expression, with 0–4 molecules of *Xist* per dataset, none in the relevant cell type.

For comparison, we also quantified the amount of *Tsix*. This gene is not explicitly annotated in the reference. However, *LOC134484082* is a reasonable candidate: this transcript is antisense to *Xist*, and its longest exon almost completely covers exon 1 of *Xist*. Interestingly, this transcript is absent in the mRatBN7.2/rn7 assembly, which annotates *Xist* as *LOC100911498* or transcript NR_132635.1. In that assembly, the putative *Tsix* (*LOC680227*) is missing the portion of the 3’ exon that covers the *Xist* promoter, previously reported by Shevchenko et al. [96] in their study of rodent *Xist* conservation.

Despite being based on the Rnor6 assembly, the Arduini et al. genome (acquired from GEO at GSE280111) did not contain *LOC100911498* or *LOC134484082*, but did contain putative *Tsix* (*LOC680227*), annotated as a protein-coding gene. We acquired count data from the Broad Single Cell Portal (SCP2828). The released dataset contained modest amounts of *Tsix* (9–44 UMIs per LV sample). This is somewhat lower than the *kallisto | bustools* quantification (53–204 UMIs), likely due to a combination of alignment strategy differences, filtering, and truncation of an exon.

##### Cynomolgus: Qu

A single-cell atlas of the cynomolgus monkey (*Macaca fascicularis*) released heart transcriptomic data. Although XCI has been studied in *M. fascicularis* [97, 98], the standard reference genomes do not contain *XIST*. The human *XIST* is fairly similar to the *Macaca mulatta* long non-coding RNA *LOC106995245* (cover 30%, identity 92.3% relative to human transcript variant 1), suggesting it as a reasonable candidate for the *M. mulatta XIST* [99]. We used Liftoff [100] to map the RefSeq version of the *M. mulatta* genome Mmul_10/rheMac10 to the Ensembl *M. fascicularis* assembly Macaca_fascicularis_6.0, localizing the putative gene to X:67,721,448–67,753,629 in the latter genome. After extracting and harmonizing [101] the *LOC106995245* reference, we concatenated it to Macaca_fascicularis_6.0 and generated a pseudoalignment reference using *kallisto | bustools* 0.28.2 [95], using kb ref in the nac configuration.

We pseudoaligned the male heart, male liver, and female liver samples released by Qu et al. using kb count in the nac configuration and inspected the unfiltered total count matrices to characterize the *LOC106995245* content of the sequencing libraries [102]. The female and male liver samples had 11,104 and 187 UMIs of *LOC106995245*, respectively, supporting the hypothesis that *LOC106995245* is the sex-specific gene *XIST*. The heart sample had a single UMI of *LOC106995245*.

The Qu et al. atlas does not explicitly report the detection of pseudo-glia, and it is conceivable that this rare cell type could simply be absent in this particular dataset. However, clustering the *scVI* embedding of bustools-filtered data yields some 300 cells substantially enriched in the pseudo-glial markers, but without any detectable *LOC106995245* [91, 92], suggesting that the population is present but does not express *XIST*.

##### Pig: Miao, Mendelson, Wang

At least four single-cell pig heart studies are available [103–106]. Three report small but distinct neuronal populations, marked by *Scn7a* [103, 106] and by *Nrxn1* [104]. In addition, a recent cross-tissue atlas sequenced heart samples from a mixture of three male pigs [107], albeit without explicitly reporting pseudo-glia (Figure S2 of [107]).

These studies’ disclosed raw data matrices do not contain either *Xist* or *LOC102165344*, the gene’s identifier in the 2011 Sscrofa10.2 assembly. Neither identifier is listed in the standard 2017 genome Sscrofa11.1. Porcine *Xist* has been annotated as locus KC149530 [108] and is known to be functional [108] in spite of nontrivial differences from the human gene [48], but characterization of its presence or absence in male porcine heart cell populations requires more comprehensive transcriptome build.

We used Liftoff [100] to map the RefSeq version of the *S. scrofa* genome Sscrofa10.2 to the Ensembl assembly Sscrofa11.1, localizing the putative gene to X:59,277,353–59,310,014 in the latter genome. After extracting and harmonizing [101] the *LOC102165344* reference, we concatenated it to Sscrofa11.1 and generated a pseudoalignment reference using *kallisto | bustools* 0.28.2 [95], using kb ref in the nac configuration.

Pig sexes were not reported in the Nakada and Wei studies [103, 105], leading us to reanalyze the remaining three studies. The Miao and Mendelson data were generated using the standard 10X Genomics 3’ v3 technology. The single-nucleus data in the Wang atlas were generated using an MGI DNBelab C4 instrument, which uses two 10-base barcodes selected from a common whitelist and 10-base UMIs. The read structure and whitelists are disclosed by the manufacturer (DNBelabC4_scRNA_readStructure.json in [109]); we manually generated the corresponding 20-base cell barcode whitelist for *kallisto | bustools*. From the annotation, we inferred that the read direction was forward. This inference is consistent with trial pseudoalignments (e.g., SRR17661076 produces 11 million UMIs in the forward configuration but 6 million UMIs in the reverse configuration).

We pseudoaligned all samples using kb count in the nac configuration, setting the *kallisto | bustools* parameters appropriately. We analyzed the 10X and DNBelab datasets separately to avoid embedding artifacts stemming from technology differences.

Clustering the *scVI* embedding of bustools-filtered data from the Miao and Mendelson studies yields 3,587 cells substantially enriched in the pseudo-glial markers, but with 0–3 molecules of *LOC102165344* per sample [104, 106], suggesting that the population is present but does not express *XIST*.

Clustering the *scVI* embedding of bustools-filtered data from the Wang study did not yield an identifiable pseudo-glial population (consistently with the original article), and the expression of the cell type’s markers was sparse. This may be an artifact of the isolation procedure, an incidental consequence of the low cell count (*≈* 3000 cells passing QC in the authors’ analysis), or, somewhat less plausibly, a consequence of the sequencing technology. The entire dataset contained *≈* 100 UMIs of *LOC102165344* across five samples, with no clear cell type enrichment.

None of the three studies had female samples that could serve as comparable positive controls for *Xist* expression. However, the Wang et al. atlas reports four samples generated from a single female pig (lung, blood, and two adipose samples), albeit using a single-cell protocol and the 10X chemistry (as opposed to single-nucleus and DNBelab used for the heart). Reanalyses of these data confirmed the substantial expression of *LOC102165344* in the lung and blood samples. However, the adipose samples had low expression of putative *Xist*, alongside high expression of the Y-linked genes *ZFY*, *KDM5D*, and *EIF1AY*, suggesting they were incorrectly labeled and originated from a distinct male pig.

### 4.2 Human data analysis and statistics

We used the cell type annotations and donor IDs, defined as in Section 4.1.1, to generate count sum pseudobulks using the Python package decoupler [110]. The parameters were as lenient as possible, with molecule and cell count thresholds set to zero.

Identification of cells in the pseudo-glial population was performed by string matching to cell type annotations. We extracted all cells whose cell type names included Glia, glia, Neuron, neuron, Neural, neural, or Schwann. This strategy takes the clusters at face value, and omits, e.g., plausibly glia-like cells assigned to other cell types or annotated as unknown.

*XIST* -positive pseudo-glia were defined näıvely, by identifying all cells belonging to the relevant clusters, binning them into zero- and nonzero-*XIST* categories based on expression, and pseudobulking based on donor ID and *XIST* presence. The presence or absence of a gene may be different between “processed” (e.g., anndata.X) and “raw” expression matrices (e.g., anndata.raw.X), which occurs when processing involves imputation (in the Eraslan, Selewa, and Sim datasets). We used the sparsity pattern of the raw data to split the datasets.

#### 4.2.1 Expression comparisons

Figures 1, 2, and 6a were generated by computing sample means: extracting all relevant pseudobulks (glial or non-glial) for a donor, adding the gene counts, then dividing them by the sum of the corresponding psbulk_n_cells entries, i.e., the total number of cells.

#### 4.2.2 Differential expression between male and female donors

To characterize the sex-dependent expression differences within the pseudo-glial cells, we aggregated all cells in these populations into coarser sum pseudobulks. Next, we fit a PyDESeq2 model using the donor sex and the source study as categorical covariates [111, 112]. Finally, we obtained the Wald test *p*-values and log fold differences for the female/male contrast and visualized them in Fig 4, providing three axis scales to accommodate the outsize effect sizes of sex chromosomes.

**Fig 4.**
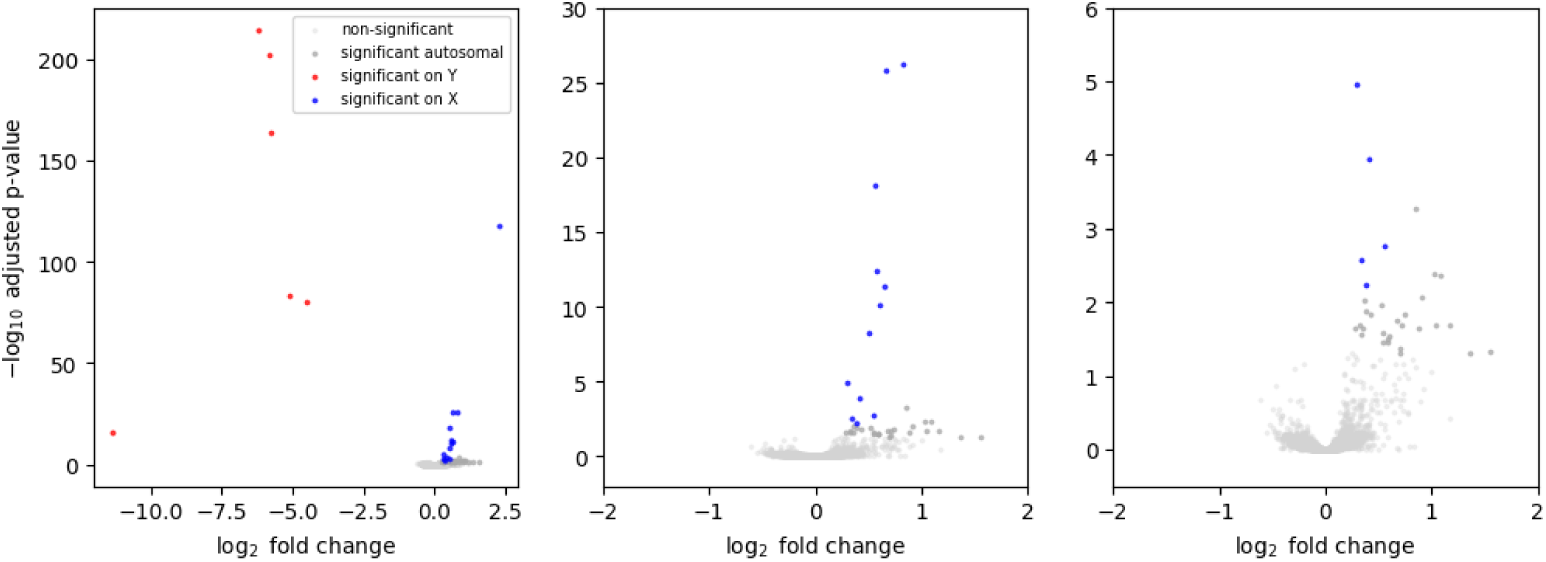
Results of differential expression testing between female and male samples, using sex and study as covariates. Genes with baseMean *<* 5 excluded from visualization. Differential expression is defined as | log_2_ FC*| >* 0.25 and adjusted *p*-value *<* 0.05.

Evidently, female samples have little to no expression of Y-chromosomal genes. However, they also have higher expression of certain X-chromosomal genes – most prominently by far, *XIST*, but also the XCI mediators *JPX* and *ZFX*. In addition, the expression of X-chromosomal genes *KDM6A*, *NLGN4X*, *AP1S2*, *EIF1AX*, and *CA5B* is 50–80% higher in the female samples, in the correct range for XCI escape (i.e., higher than in male, but not quite doubled).

It is curious to note that the sex analysis is analogous to a cross-study investigation very recently released by Gao et al. [34]. The volcano plots shown in Fig 4 partially reproduce Supplemental Figure 15a of that preprint. In addition, the results of this differential expression test immediately imply the key finding of the current study: the expression of *XIST* in the pseudo-glia (the “neuronal” population) is approximately fourfold lower in males than in females, about half an order of magnitude (Fig 1). This is in stark contrast to other cell types, where the male expression of *XIST* is far more sparse (log_2_ fold change below *−*5, and frequently even lower).

For the sake of completeness, we note that the analysis of epicardial cells (*ITLN1* /*MSLN*) in Supplemental Figure 9a of Gao et al. reports sex-specific enrichment of *XIST* quantitatively similar to the pseudo-glia, with log_2_ fold change *≈ −*2. However, the cells expressing these markers (typically annotated as “mesothelial”) do not appear to exhibit systematic male *XIST* expression [23, 31–33, 40]. Therefore, the reported enrichment of *XIST* in male epicardial cells may be an artifact of the drastic differences in this cell type’s representation between studies (7% of cells in [32], 0.15% in [23], 0.1% in [31], and not readily detectable in [33, 41]). This imbalance may lead to suboptimal clustering or to inaccurate estimation of pseudobulk “size factors,” either of which can lead to miscalibration of differential expression testing.

#### 4.2.3 Differential expression between *XIST* -positive and -negative cells

Ideally, the study of *XIST* in the pseudo-glial population should fully characterize the impact of its presence or absence on the molecular makeup and phenotype of the cell population, its niche, the tissue, and the organism. Thus far, there is no evidence that this expression pattern occurs outside humans (Section 4.1.2), precluding *in vivo* study. It is possible that part of the context may be recapitulated with an appropriately constructed tissue model. A recent study reported that a heart organoid model produces a small population of conductance cells [113] that resemble embryonic Schwann and neural cell progenitors [114]. However, this population’s markers are not expressed in adult pseudo-glia, so the generation of a model that recapitulates the adult heart remains an outstanding problem.

In the absence of an experimental system that can be perturbed (e.g., by knocking out *XIST*), it is possible to approximate regulatory relationships between genes using endogenous patterns of co-expression. This approach falls under the umbrella of gene regulatory network (GRN) inference. Although this is a thriving field with numerous tools, the problem of GRN inference is far from solved [115–117], and applying this class of algorithms to a rare cell type scattered across hundreds of donor samples collected under different conditions is unlikely to be informative.

In lieu of attempting to discover genome-wide networks, we restrict ourselves to describing the immediate impacts of *XIST* presence or absence. This classification is necessarily imperfect. Due to contamination, cells with no *XIST* may be reported as having *XIST*. Conversely, due to the limitations of cDNA library construction and sequencing, some molecules may not be captured. The background contamination – if comparable to that in the heart overall (tentatively, the yellow triangles in Fig 1) – is unlikely to be more than a minuscule fraction of the total. The issue of false negatives is more challenging to bypass. However, the misassignment of *XIST* -positive cells as *XIST* -negative attenuates, rather than increases differences between the categories, so this approach is conservative.

Bearing these limitations in mind, we investigated the differential expression of genes between the *XIST* -positive and -negative cells, applying PyDESeq2 to pseudobulks. As we are interested in isolating the effects of intra-individual variation, we encoded the donor ID as a categorical covariate alongside *XIST* status. This results in a relatively complex problem, with 226 covariates for the 225 male donors with *XIST* expression and 90 covariates for the 89 corresponding female donors. We fit the model using the intersection of genes identified in all datasets. We used a PyDESeq2 Wald test to compute *p*-values, then filtered the results further for genes with baseMean *>* 5. As the *p*-values are adjusted based on the entire model, the values shown in Fig 3 are conservative. We note this filtering is rather stringent, as it discards some 78% of the 10,391 genes intersecting between datasets. Although it is likely that some of the omitted genes are important, the small sample sizes (61% of the male pseudobulks have fewer than fifty cells; 16% have fewer than ten) produce an considerable risk of false positives for low-expression genes.

Some reasonable artifactual explanations for the differential expression patterns observed in Fig 3 involve the diffusion of abundant myogenic transcripts into droplets containing pseudo-glia. In other words, the enrichment of background genes may take place due to insufficiently accounted-for “size” effects, which may occur through technical or combination technical and biological mechanisms. In the first scenario, *XIST* -negative populations incidentally have fewer overall sequencing primers, and capture fewer endogenous and exogenous transcripts. As shown in Fig 5, this intuition holds for female samples but does not hold for male samples, which tend to have somewhat *less* RNA.

**Fig 5.**
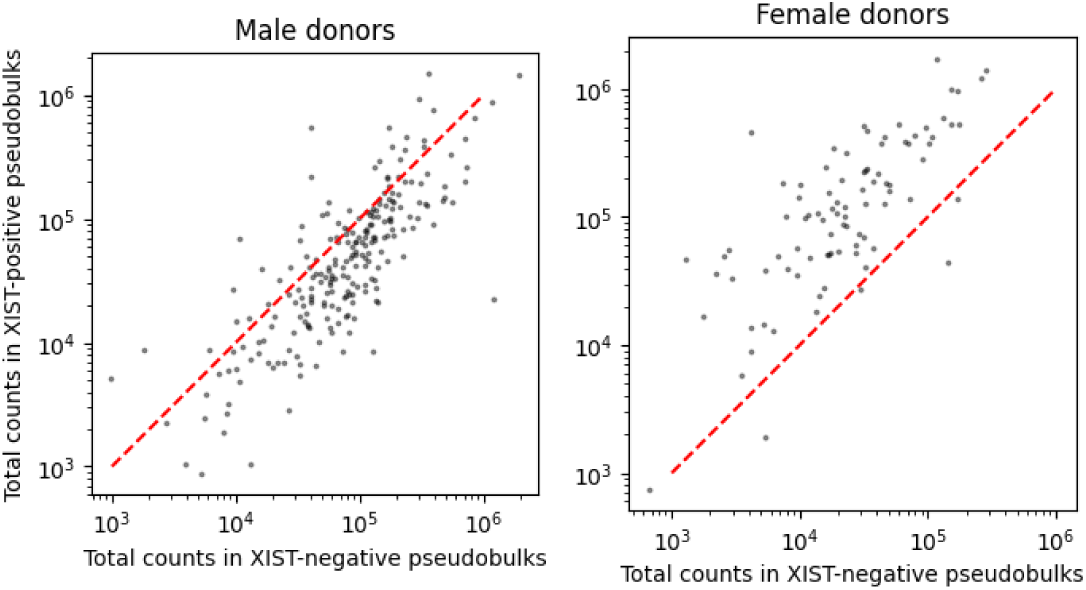
Total molecule counts in the pseudobulks used to evaluate differential expression between *XIST* -positive and -negative cells (points: donor pseudobulks; lines: identity).

In the second scenario, *XIST* -positive populations systematically have lower overall endogenous genome-wide expression (presumably due to *XIST* -mediated repression, incidentally consistent with Fig 5), and relatively more of the cDNA library is sourced from the contaminating transcripts. In both scenarios, one would expect *XIST* -positive populations to have *higher* rather than lower expression of ubiquitous muscle genes, the opposite of what is observed. Although it is challenging to make definitive conclusions based on summary data collected using heterogeneous procedures, the comparison of paired samples is rather more suggestive of endogenous suppression than exogenous contamination.

#### 4.2.4 *XIST* relationship to age

To characterize the age-dependent expression variability within the pseudo-glial cells, we followed the procedure in Section 4.2.2 to construct coarse pseudobulks, subset the data to non-fetal males with age annotations, then fit a PyDESeq2 model using the source study and the age as covariates. As elsewhere, when only coarse bins were available, we used the bin midpoint as an imperfect proxy for the true age.

First, we used a model which directly used the age as a continuous covariate. The fit yielded an effect size of 0.007, with a *p*-value of 0.05. This effect is small and dubiously significant. Next, we used a model which used the logarithm of the age as the covariate. This analysis yielded an effect size of 0.27, with a *p*-value of 0.01. In contrast, the genes with the lowest *p*-values in the latter analysis, filtering for those with baseMean *>* 10, were *PPARG* (effect size 0.89 and *p*_adj_ = 5.9 *×* 10*^−^*^3^) and *FSTL3* (effect size 0.71 and *p*_adj_ = 7.8 *×* 10*^−^*^3^), both of which are known to serve roles in heart failure pathology [118–120]. We note that the targeted inspection of *XIST* uses the nominal *p*, whereas the untargeted inspection differential expression tables uses the *p*-value adjusted for multiple comparisons. Although it may be the case that *XIST* slightly increases in adolescence, there does not appear to be strong evidence for a continuing increase in expression during adulthood, whether considering the sample mean (Fig 6a), the PyDESeq2 “size-normalized” counts (Fig 6b, top), or the latter with additional correction for the batch covariates (Fig 6b, bottom). In contrast, the age dependence of *PPARG* and *FSTL3* is somewhat less equivocal, if slight. However, these results are somewhat challenging to interpret without appropriately accounting for the uncertainty represented by binned age metadata.

**Fig 6.**
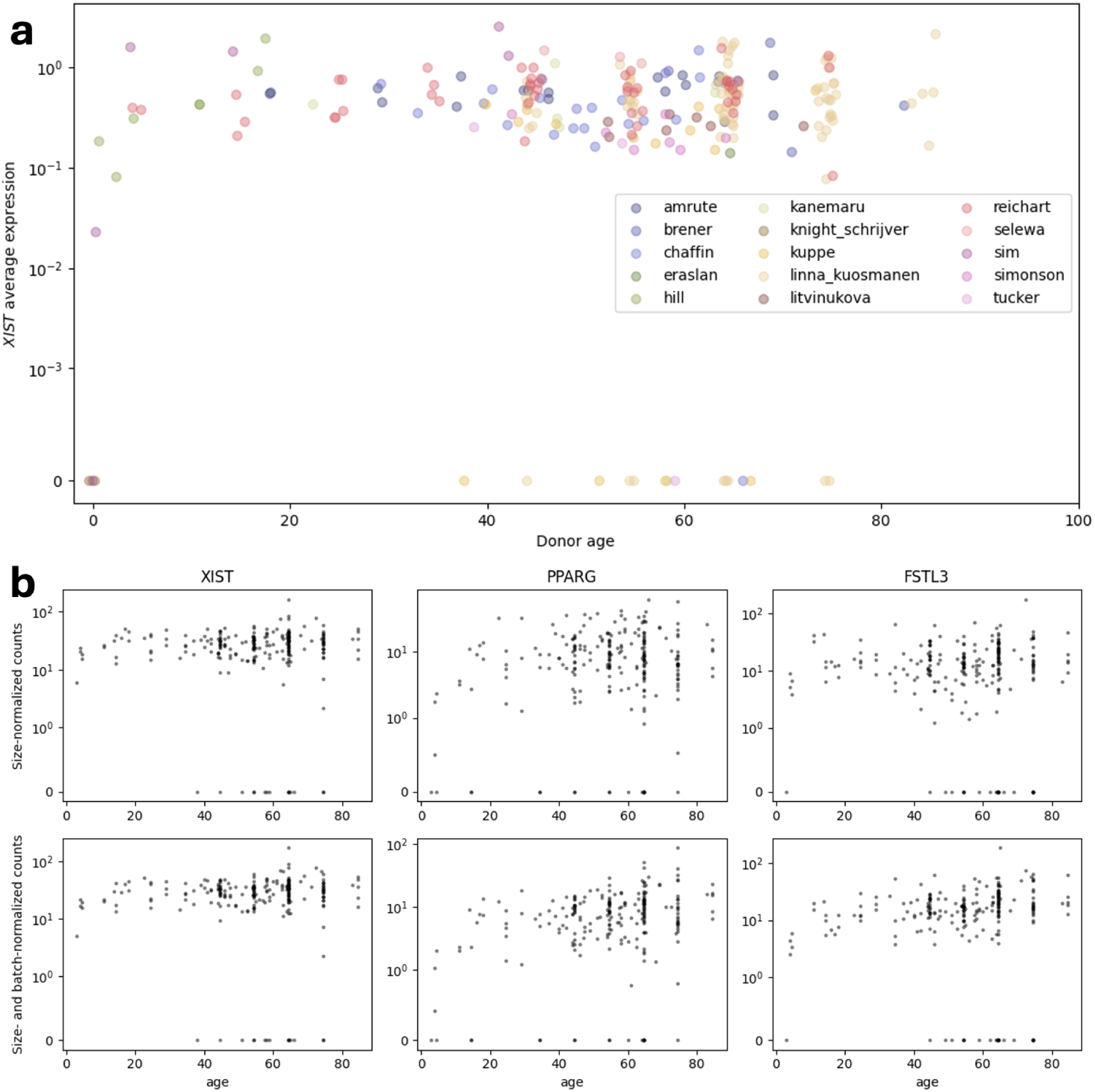
Age dependence of cardiac gene expression. **a.** Average expression of *XIST* in pseudo-glia in male donors across ages. Colors denote study of origin. All fetal samples are given age zero. Donors with no age information omitted. Jitter added. Data with binned ages (e.g., 20–29 years) are visualized at the bin midpoint. To accommodate zero values, the region [0, 10*^−^*^3^] is linear. **b.** Expression of *XIST*, *PPARG*, and *FSTL3* in pseudo-glia in male donors. Data with binned ages (e.g., 20–29 years) are visualized at the bin midpoint. Top: size-normalized pseudobulk counts. Bottom: size-normalized pseudobulk residuals, after correction for batch variables, inferred from a batch/log-age fit. To accommodate zero values, the region [0, 1] is linear.

**Fig 7.**
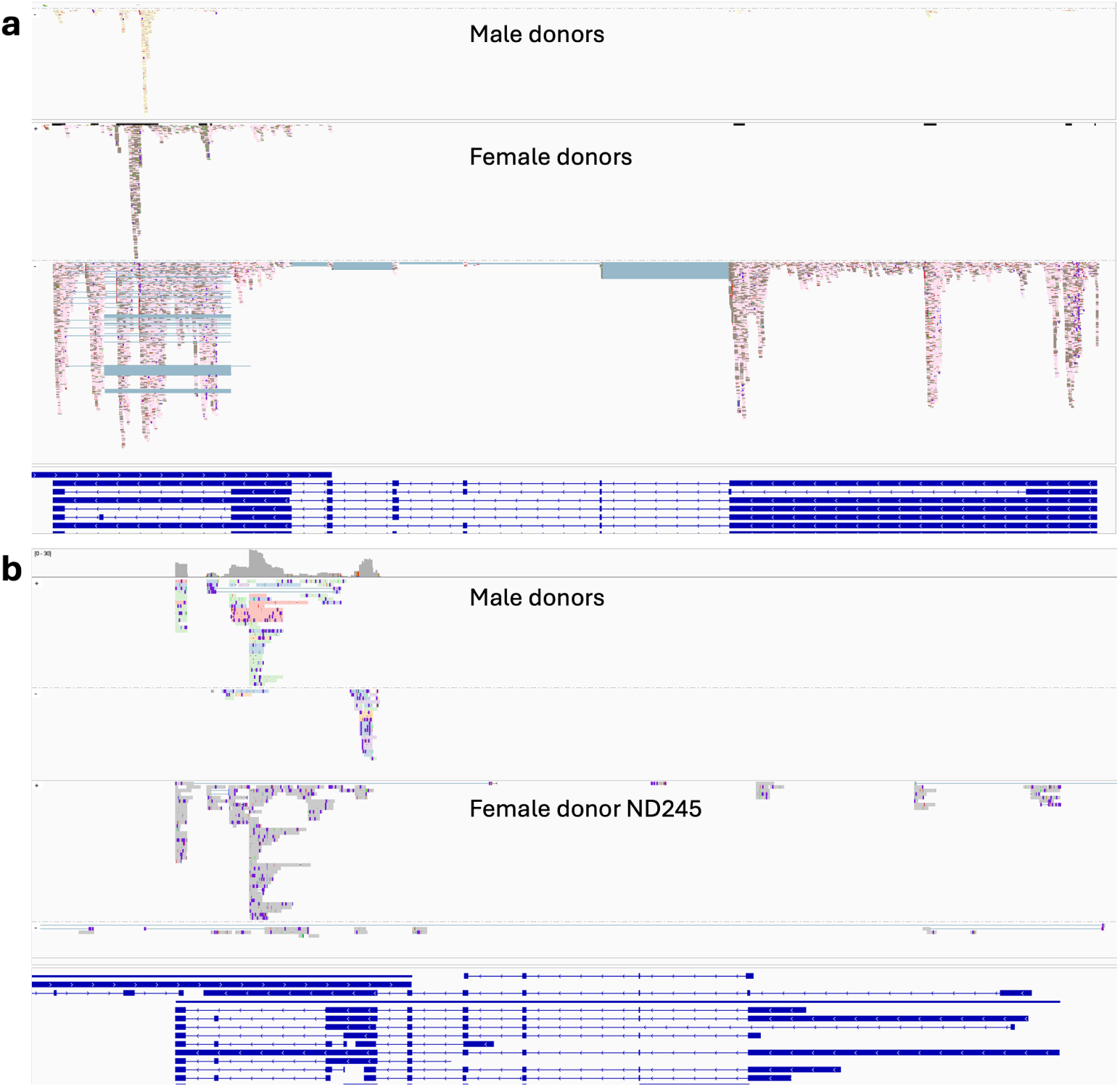
Read coverage and distribution of *XIST* transcripts. **a.** Read alignments of Larson and Chin [44] septum samples. Top panel: male donors (color-coding: donor identity; dark blue lines: splice junctions; reads are split by strand, with positive at the top). Middle panel: female donors (conventions as in top panel). Bottom panel: Representative RefSeq transcripts in the region. **b.** Read alignments of Pan et al. [45], restricted to pseudo-glial samples (conventions as in **a**, with the negative strand at the top of each subpanel).

#### 4.2.5 *XIST* relationship to pathology

Given the abundance of pathological samples in several publicly available studies, as well as the lack of evidence regarding the protective or pathological roles of *XIST*, we selected the largest datasets for exploratory analysis of the relationship between *XIST* expression and heart failure phenotypes.

##### Chaffin

The dataset released by Chaffin et al. had data for healthy controls as well as hypertrophic and dilated cardiomyopathy (HCM and DCM) patients. We separately compared *XIST* expression between the pathology classes and healthy controls. A PyDESeq2 Wald test yielded higher expression in the HCM/control comparison (effect size 0.88 and *p* = 1.3 *×* 10*^−^*^3^) and in the DCM/control comparison (effect size 0.98 and *p* = 1.3 *×* 10*^−^*^3^), with similar results under models incorporating age or the logarithm of age alongside the pathology.

##### Reichart

The dataset released by Reichart et al. had data for healthy controls as well as several classes of cardiomyopathy. We compared *XIST* expression between the largest classes (DCM and arrhythmogenic right ventricular cardiomyopathy, ARVCM) and healthy controls. A PyDESeq2 Wald test yielded higher expression in the DCM/control comparison (effect size 1.0 and *p* = 5.9 *×* 10*^−^*^4^) and in the ARVCM/control comparison (effect size 1.3 and *p* = 9.9 *×* 10*^−^*^4^) with similar results under models incorporating age or the logarithm of age alongside the pathology.

##### Linna–Kuosamanen

The dataset released by Linna–Kuosamanen et al. had patients with a variety of conditions; all patients had some form of valve disease. In addition, the released AnnData files annotated numerous donors as having atrial fibrillation (AF), in conflict with the metadata: for instance, of the fourteen non-control Periheart donors, zero are reported as having AF in the article, but four are reported as having AF in the dataset. As the original publication treated valvular disease patients as “controls,” we binned the samples accordingly, comparing patients with comorbidities to valve disease-only patients. A PyDESeq2 Wald test yielded slightly higher expression in the comorbidity class (effect size 0.27 and *p* = 0.035), with similar results under models incorporating age or the logarithm of age alongside the pathology.

##### Kuppe

The dataset released by Kuppe et al. compared twelve myocardial infarction (MCI) patients to three controls. A PyDESeq2 Wald test yielded slightly, but non-significantly lower expression in the MCI class (effect size *−*0.55 and *p* = 0.55), with similar results under models incorporating age or the logarithm of age alongside the pathology.

##### Simonson

The dataset released by Simonson et al. compared three ischemic cardiomyopathy (ICM) patients to four controls. A PyDESeq2 Wald test yielded slightly, but non-significantly lower expression in the ICM class (effect size *−*0.95 and *p* = 0.18), with similar results under models incorporating age or the logarithm of age alongside the pathology.

##### Summary

Evidently, the Chaffin, Reichart, and Linna–Kuosamanen datasets suggest stronger *XIST* expression in more pathological samples, whereas the others are not informative. As suggested in Section 3, reanalysis would be necessary to draw quantitative conclusions. For instance, plausible pseudo-glia may have been removed in the authors’ QC pipelines, or incidentally assigned to other cell types, biasing the expression profiles, and a reanalysis may recover them. However, it is unlikely that even such follow-up would be conclusive: subtle details may differ by age, sex, chamber, pathology, its severity, and perhaps subpopulation. Given these caveats, as well as the rarity of the cell type, it appears likely that the pathological role, if any, can only be determined with thorough characterization of its function in healthy tissues, followed by the extensive and targeted collection of new data.

### 4.3 Short read analysis

Read alignment files were acquired from the Sequence Read Archive depositions accompanying the GEO accession GSE161921 [44]. Specifically, we acquired the sorted BAM files generated by Cell Ranger alignments of data for male donors 2879 and 2880, as well as female donors 2867 and 2884. The alignments were filtered to retain only the reads overlapping *XIST* using the samtools 1.20 command view for the region X:73818656-73854714, then filtered further to include only reads with the GN tag [121].

This procedure retains all reads that would normally be quantified, but omits, e.g., uncharacterized transcripts. The resulting subsets were visualized using the Integrative Genomics Viewer [122] with reference to the GRCh38/hg38 genome.

### 4.4 Long read analysis

A recent study by Pan et al. [45] constructed 10X cDNA libraries from isolated heart nuclei, then sequenced them using long-read nanopore (ONT PromethION) sequencing, aiming to characterize the isoform diversity of the healthy and pathological human heart. We cross-referenced the findings from the 10X/Illumina datasets from other studies against the Pan et al. data, but did not analyze them jointly.

Gene quantification results were obtained from GEO accession GSE288222. Sample-and cell-level metadata were obtained from Tables S1 and S8 of the original publication, with 28,534 out of 59,488 barcodes having cell type annotations.

The pseudo-glial population was only reported in 4/6 female samples and 5/6 male samples. Three male samples had no *XIST* expression in this cell type, and one had negligible expression at a level of *≈* 0.01 UMIs per barcode (as opposed to 0.9–3.1 UMIs per barcode in female samples). We hypothesized that this result is a consequence of the alignment strategy: as the reads reported in Section 4.3 are predominantly localized to the 3’ exon of *XIST*, an unstranded alignment would be unable to disambiguate *XIST* reads from antisense *TSIX* reads. The authors do not explicitly report the alignment setting, which is chosen automatically as part of the *Minimap2* pipeline [123], but one corroborating line of evidence is the amount of *TTN-AS1*, a transcript antisense to titin. Previous heart atlases (e.g., [23, 31]) report considerable amounts of *TTN-AS1* in cardiomyocytes, but the counts are relatively low in the Pan et al. data. However, it is challenging to judge the discrepancies: *TTN-AS1* reported in short-read atlases may itself be a *TTN* -derived antisense artifact that the nanopore protocol happens to avoid. In addition, the discrepancies may result from the omission or misassignment of cells in the rare population due to pre-processing choices.

We obtained six male datasets and one female dataset (ND245) from the Sequence Read Archive and split the nanopore reads into technical/biological read pairs using the *scNanoGPS* utility Scanner [124] per the authors’ procedure. This produced a set of reads in the reverse configuration, starting with a poly(T) tail (as opposed to forward-configuration reads conventionally produced by the 10X protocol).

At this juncture, we diverged from the Pan et al. procedure. We generated a *kallisto | bustools* index from the human genome (10X Genomics, 2024-A version) using *k* = 63, then used the long-read kb count pipeline [125] in the nac configuration, assuming reverse strandedness, to pseudoalign the reads.

Clustering the *scVI* embedding of knee plot-filtered data yields 364 cells substantially enriched in the pseudo-glial markers (10–136 per male sample).

Pseudo-glia show average male *XIST* expression between 0.06 and 0.5 UMIs per barcode (1–29 UMIs per pseudobulk), whereas other cells (excluding empty drops) show expression below 0.008 UMIs per barcode, consistently with Figure 1. As expected, the female sample shows high *XIST* across the cellular landscape (*≈* 1.4 UMIs per barcode in pseudo-glia, 1.9 otherwise). Curiously, *TTN-AS1* is present in relatively high amounts (tens of thousands of UMIs, vs. dozens of UMIs in disclosed data), suggesting that the antisense titin cDNA are either derived from physiological molecules or pre-fragmentation artifacts.

In the interest of being exhaustive, we tested the differential expression of *XIST* between 4 heart failure and 2 non-diseased samples, as in Section 4.2.5. As may be expected from the small sample size, the comparison was inconclusive (effect size *−*0.41 and *p* = 0.64).

In principle, long-read sequencing can be used to elucidate transcript identities better than short-read sequencing. To check the transcript structures, we identified barcode/UMI pairs for reads mapping to *XIST* and *TSIX*, extracted the corresponding entries from the biological read files, trimmed the poly(T) sequences, and remapped them to a section of the genome immediately surrounding this gene pair using *Minimap2* [123].

## Data availability

No new experimental data were generated for this study. All datasets investigated here are available at, and were obtained from, CZI CELL*×*GENE, Gene Expression Omnibus, the Broad Single Cell Portal, the Human Cell Atlas Data Explorer, or the Sequence Read Archive. The sole exception is the set of annotated data originally released on the Single Cell Portal by Brener et al. [24], which was obtained prior to its removal from the database. The dataset released on GEO contains data for the same cells and genes, and information regarding donor identities and various quality control metrics, but does not contain cluster identities; to our knowledge, there are no publicly available resources that continue to host this dataset. The details of the data acquisition and curation process are given in Section 4.1.

The Python notebooks used to perform the analyses in this study are hosted at https://github.com/Fauna-Bio/GDG_2025. The processed data (differential expression tables, *XIST* -aligning reads, count matrices, pseudobulks used for statistical tests, and genome reference entries) are hosted on Zenodo [126]. The intermediate data (split long reads, pseudoalignments) are available on request.

The differential expression analyses used PyDESeq2 0.4.8 [111]. The visualization of reads used the Integrative Genomics Viewer [122]. The pseudoalignment of rat data used *kallisto | bustools* 0.28.2 with the GRCr8 genome. The pseudoalignment of cynomolgus data used *kallisto | bustools* 0.28.2 with the Macaca_fascicularis_6.0 genome, augmented with the putative *M. mulatta XIST*. The pseudoalignment of pig data used *kallisto | bustools* 0.28.2 with the Sscrofa11.1 genome, augmented with the putative Sscrofa10.2 *XIST*. The pseudoalignment of long-read heart data used *kallisto | bustools* 0.29.5 with the 2024 version of the GRCh38 genome released by 10X Genomics. The alignment of putative *XIST* long reads used *Minimap2*.

## Acknowledgments

L.G. is a co-founder and CTO of Fauna Bio. G.G. and N.D. are employees of Fauna Bio. The authors would like to acknowledge Tara Chari, Sina Booeshaghi, Prashant Bhat, Phil McNamara, Evelyn Tran, Alan Cohen, Natalie DeForest, and Alex Burr for their assistance in the preparation of this manuscript. The visualizations in this manuscript use the LaCroixColoR palette by Dave Armitage and Johannes Bjork.

